# Mapping Disease Transitions from Premalignant, Asymptomatic to Advanced Myeloma through Integrative Epigenomic and Transcriptional Analyses

**DOI:** 10.64898/2026.06.03.729851

**Authors:** Aleksandra Kurowska, Azari Bantan, Raghad Shuwaikan, Nahia Gomez-Echarte, Itziar Cenzano, Paula Aguirre-Ruiz, Estíbaliz Miranda, Leire Garate, Amaia Vilas-Zornoza, Patxi San-Martin, Marta Larrayoz, Beñat Ariceta, Ane Amundarain, Daniel Soto Ortiz, Markos Tsitsianopoulos, Argyris Papantonis, María José Calasanz, José Ignacio Martin-Subero, Vincenzo Lagani, Jesper Tegner, Jose A. Martinez-Climent, Mikel Hernaez, Nuria Planell, Xabier Agirre, Felipe Prosper, David Gomez-Cabrero

**Affiliations:** Division of Biomedical Sciences, King Abdullah University of Science and Technology KAUST, Thuwal, Saudi Arabia; Computational Biology Program, Centre for Applied Medical Research (CIMA), Instituto de Investigaciones Sanitarias de Navarra (IdiSNA), Cancer Center Clinica Universidad de Navarra (CCUN), Pamplona, Spain; Hematology and Oncology Program, Centre for Applied Medical Research (CIMA), Instituto de Investigaciones Sanitarias de Navarra (IdiSNA), Cancer Center Clinica Universidad de Navarra (CCUN), Pamplona, Spain; Centro de Investigacion Biomedica en Red de Cancer (CIBERONC), Madrid, Spain; Institute of Pathology, University Medical Center Göttingen, Göttingen, Germany; Hematological Diseases Laboratory, CIMA LAB Diagnostics, Clinica Universidad de Navarra, 31008 Pamplona, Spain; Institució Catalana de Recerca i Estudis Avançats (ICREA), Barcelona, Spain; Institut d’Investigacions Biomèdiques August Pi i Sunyer (IDIBAPS), Barcelona, Spain; Institute of Chemical Biology, Ilia State University, Tbilisi, Georgia; Center for Data Science and Artificial Intelligence (DATAI), University of Navarra, Pamplona, Spain; Hematology and Cell Therapy Department. Clinica Universidad de Navarra, IdiSNA, Pamplona, Spain

**Author notes:** contributed equally (Aleksandra Kurowska, Azari Bantan, Raghad Shuwaikan). contributed equally (Nahia Gomez-Echarte, Itziar Cenzano).

## Abstract

Multiple myeloma (MM) evolves from asymptomatic precursor conditions through progressive genetic and epigenetic remodeling, yet the regulatory mechanisms driving the development toward more active stages of the disease remain poorly understood. Here, we integrated bulk-based paired chromatin accessibility and activation, and transcriptomic profiling across disease stages to map regulatory remodeling during myeloma development. We identified a progressive increase in chromatin accessibility; furthermore, this epigenetic reconfiguration is accompanied by a stage-dependent shift from promoter-centered regulation in precursor states toward enhancer-dominated transcriptional control in active MM. Motif enrichment and regulatory network analyses identified both established and previously underappreciated transcription factors (TFs), including members of the IRF, MEF2, and FOX families, associated with disease-stage–specific transcriptional programs. Among these, MEF2D and FOXK2 emerged as candidate regulators of pathways involved in cell survival and chemotaxis. Functional perturbation demonstrated that MEF2D depletion markedly impaired MM cell viability, whereas inhibition of either MEF2D or FOXK2 reduced chemotactic migration. Together, these findings provide a stage-resolved framework of epigenetic and transcriptional remodeling across myeloma development, revealing regulatory programs established in precursor conditions and progressively reinforced during disease evolution, while identifying candidate transcriptional dependencies with potential biological and therapeutic relevance.

## Introduction

Multiple myeloma (MM) is a heterogeneous hematological malignancy characterized by the clonal expansion of plasma cells (PCs) within the bone marrow (BM), leading to immune dysfunction and end-organ damage. Despite therapeutic advances, MM remains incurable due to recurrent relapse and the emergence of refractory disease^1^.

MM develops along a clinical continuum from asymptomatic precursor stages to active disease. Monoclonal gammopathy of undetermined significance (MGUS) is defined by a clonal BMPC population comprising less than 10% of BM cells and carries an annual risk of progression of approximately 1%, whereas smoldering MM (SMM), characterized by a higher tumor burden, is associated with a substantially increased risk of progression (approximately 10% per year)^1,2^. Disease progression is accompanied by the accumulation of genetic cytogenetic abnormalities^3^ and widespread epigenetic dysregulation^4^. In particular, MM cells are characterized by *de novo* chromatin activation and changes in DNA methylation, collectively reshaping gene regulatory networks to sustain malignant cell survival and proliferation^4–8^.

Previous multi-epigenomic studies have demonstrated extensive activation of regulatory elements in neoplastic PCs leading to transcriptional rewiring through enhancer-promoter interactions^9,10^. Several transcription factors (TFs), including MAF, MYC, IRF4, IRF2, XBP1, and PRDM1, have been identified as central regulators of MM biology^11–15^. More recently, integrative analyses combining RNA-seq and ATAC-seq across cytogenetic subgroups have uncovered both known and novel regulatory factors, including members of the IRF and MEF2 families that contribute to enhancer activation^16^. However, many of these studies have focused on symptomatic MM or treated patient cohorts as homogeneous groups, limiting the ability to resolve regulatory heterogeneity and its relationship to disease stages.

To address this gap, we integrate chromatin accessibility, histone acetylation (H3K27ac), and transcriptomic profiles from primary PCs spanning healthy donors and MM disease stages (MGUS, SMM, and MM). We identify stage-associated remodeling of the regulatory landscape, characterized by a widespread gain of chromatin accessibility and the emergence of active regulatory elements from previously low-signal regions. Notably, we observe a shift from promoter-centered regulation in precursor stages toward increased reliance on enhancer-associated regulatory interactions in MM. This rewiring is accompanied by the redistribution of TF activity within newly accessible regulatory regions. Among these TFs, MEF2D and FOXK2 emerge as candidate regulators linked to enhancer-driven transcriptional programs and adverse clinical outcomes, while their inhibition reduces cell viability and chemotaxis.

Together, our results provide a high-resolution map of epigenetic remodeling across myeloma disease stages and define the regulatory architecture underlying malignant PC states. These findings offer a framework for understanding the epigenetic basis of transcriptional heterogeneity in MM and highlight candidate regulatory dependencies with potential clinical relevance.

## Results

### Cumulative epigenetic changes in multiple myeloma development

To investigate regulatory mechanisms underlying MM development, we isolated BM–derived CD138+ PCs from a total of 197 patients spanning disease stages, including MGUS (*n*=15), SMM (*n*=30), and newly diagnosed MM (NDMM; *n*=145), together with seven healthy controls (HC) (**Fig. 1A, Supplementary Table 1**). For each sample, we aimed to generate matched profiles of chromatin accessibility (ATAC-seq, n=185), histone acetylation (H3K27ac ChIP-seq, n=128), and gene expression (RNA-seq, n=197), enabling integrative analyses of regulatory changes across disease stages (**Fig. 1A**). The main myeloma-associated cytogenetic alterations were analyzed in our cohort of patients (**Fig. 1B**).

**Figure 1.**
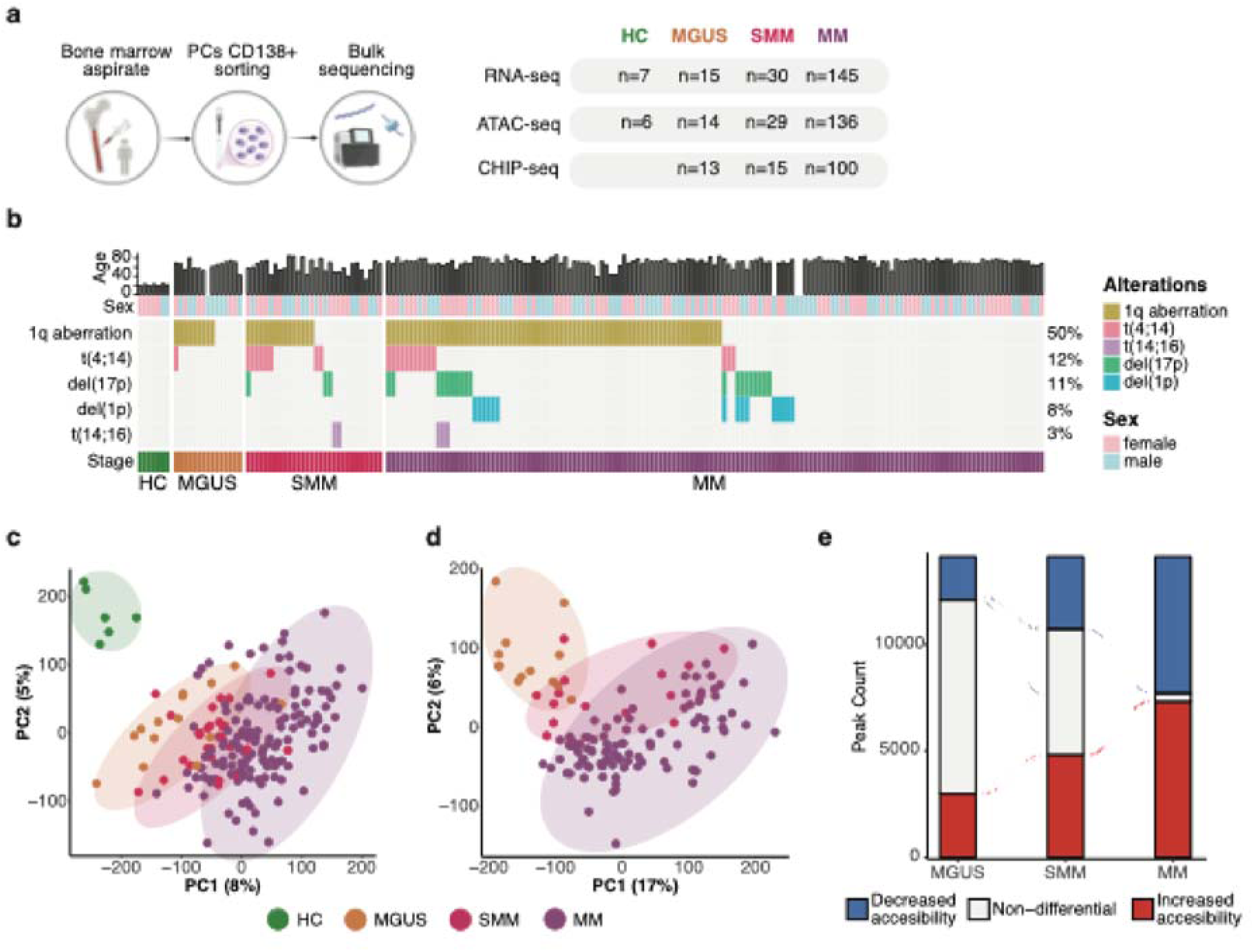
Chromatin profiling reveals progressive remodeling of regulatory chromatin during MM evolution. a. Study design and cohort overview. BM aspirates were obtained from HC and patients with MGUS, SMM and MM. CD138^+^ PCs were isolated and subjected to bulk profiling of transcriptome (RNA-seq), chromatin accessibility (ATAC-seq), and H3K27ac activity (ChIP-seq). Numbers of profiled samples per assay and disease stage are indicated within the panel. b. Summary of age, sex, disease stage, and recurrent cytogenetic abnormalities detected by fluorescence *in situ* hybridization (FISH) across all analyzed samples. Percentages shown at right indicate cohort frequency of each abnormality. Each column represents one individual sample. c. PCA based on ATAC-seq signal across 185 samples. Each point represents one sample colored by disease stage (HC, MGUS, SMM, MM). Ellipses indicate group dispersion. d. PCA of H3K27ac ChIP-seq signal across all detected acetylated regions (n = 128 samples). Each point corresponds to one sample colored by disease stage. Ellipses indicate group dispersion. e. Alluvial plot showing differential OCRs identified in pairwise comparisons of MGUS vs HC, SMM vs HC, and MM vs HC, restricted to OCRs overlapping H3K27ac peaks and positively correlated with H3K27ac signal. Red indicates regions with increased accessibility, blue indicates decreased accessibility, and gray indicates non-differential OCRs.

Because MM progression from MGUS to symptomatic myeloma is accompanied by an increasing percentage of malignant PC, and because we aimed to characterize regulatory changes occurring within malignant PCs during disease development, we estimated malignant infiltration for each sample. As direct estimates were unavailable, we inferred a non-malignant PC gene set activity score for each MGUS and SMM sample, hereafter referred to as the % normal PCs (nPCs), using transcriptional profiles and a reference gene set comprising flow-cytometry markers commonly used to distinguish non-malignant to malignant PCs, including CD19, CD27, CD38, CD45, and CD81 (see Methods for details; **Supplementary Fig. 1A)**. For subsequent analyses, we assumed that HC samples consisted predominantly of non-malignant PCs, whereas MM samples consisted predominantly of malignant PCs.

To address the epigenetic changes during disease development, we first characterized chromatin accessibility across disease stages using ATAC-seq. We identified 142,136 open chromatin regions (OCRs) mapped to annotated genomic regions using the hg38 reference genome (see Methods; **Supplementary Fig. 1B**). Those OCRs lay on a continuous chromatin accessibility trajectory separating HC from malignant stages (**Fig. 1C**). Notably, 1,528 (98.2%) of the previously identified *de novo* active regions in MM relative to HC^10^ overlapped OCRs detected in our dataset (**Supplementary Fig. 1C**). Following OCR identification, we performed differential accessibility analysis. Overall, disease stage emerged as the primary source of variation in chromatin accessibility across the majority of OCRs (**Supplementary Fig. 1D**), exceeding the contribution of the cytogenetic lesions or % nPCs. We identified 28,075 differentially accessible regions (DARs) associated with disease development.

We next evaluated the extent to which chromatin accessibility was coordinated with genomic activation. To this end, we used H3K27ac ChIP-seq data, a histone mark associated with active enhancers and promoters. H3K27ac analysis identified 93,674 acetylated regions mapped to annotated genomic elements (**Supplementary Fig. 1B**). Consistent with ATAC seq profiles, PCA of H3K27ac signal showed stage dependent separation, suggesting similar changes of chromatin accessibility and histone acetylation across stages, particularly between MGUS and SMM (**Fig. 1D**). Moreover, more than half of OCRs (n=77,361, 54.43%) overlapped H3K27ac peaks and displayed significant positive correlations between accessibility and acetylation across stages (**Supplementary Fig. 1E**), highlighting that a large fraction of OCRs represents active chromatin regulatory regions. To focus on bona fide active regulatory elements, we retained those DARs positively correlated with H3K27ac signal for all downstream analyses (n=14,135).

When we analyzed changes in accessibility and activation, chromatin accessibility correlated with H3K27ac signal across MGUS, SMM, and symptomatic MM stages, with an increase in accessibility as the disease develops (**Fig. 1E, and Supplementary Table 2**) (HC vs MGUS, n=5,026 (3,002 increased, 2,024 decreased accessibility); HC vs SMM, n=8,241 (4,834 increased, 3,407 decreased accessibility); HC vs MM, n=13,745 (7,327 increased, 6,418 decreased accessibility)). Notably, previously reported *de novo* active regions in MM^10^, showed increasing overlap with more accessible OCRs across MGUS (484; 31%), SMM (616; 40%), and MM (721; 46%), also exhibiting the cumulative pattern toward MM (**Supplementary Fig. 1F**).

Together, these results indicate a cumulative increase in chromatin accessibility alterations from non-malignant to malignant stages of MM disease.

### Chromatin accessibility dynamics shape transcriptional programs during myeloma development

To determine how this epigenetic remodeling may influence transcriptional programs, we first characterized transcriptomic dynamics in the cohort using the RNA-seq data (**Fig. 1A**). PCA of gene expression (**Fig. 2A)** partially mirrors the patterns observed in epigenomic data (**Fig. 1C, 1D**). Differential expression analysis (see Methods) identified stage-dependent transcriptional shifts, characterized by cumulative gene upregulation and downregulation, consistent with the progressive chromatin remodeling observed across disease stages (**Fig. 2B, Supplementary Fig. 2A**).

**Figure 2.**
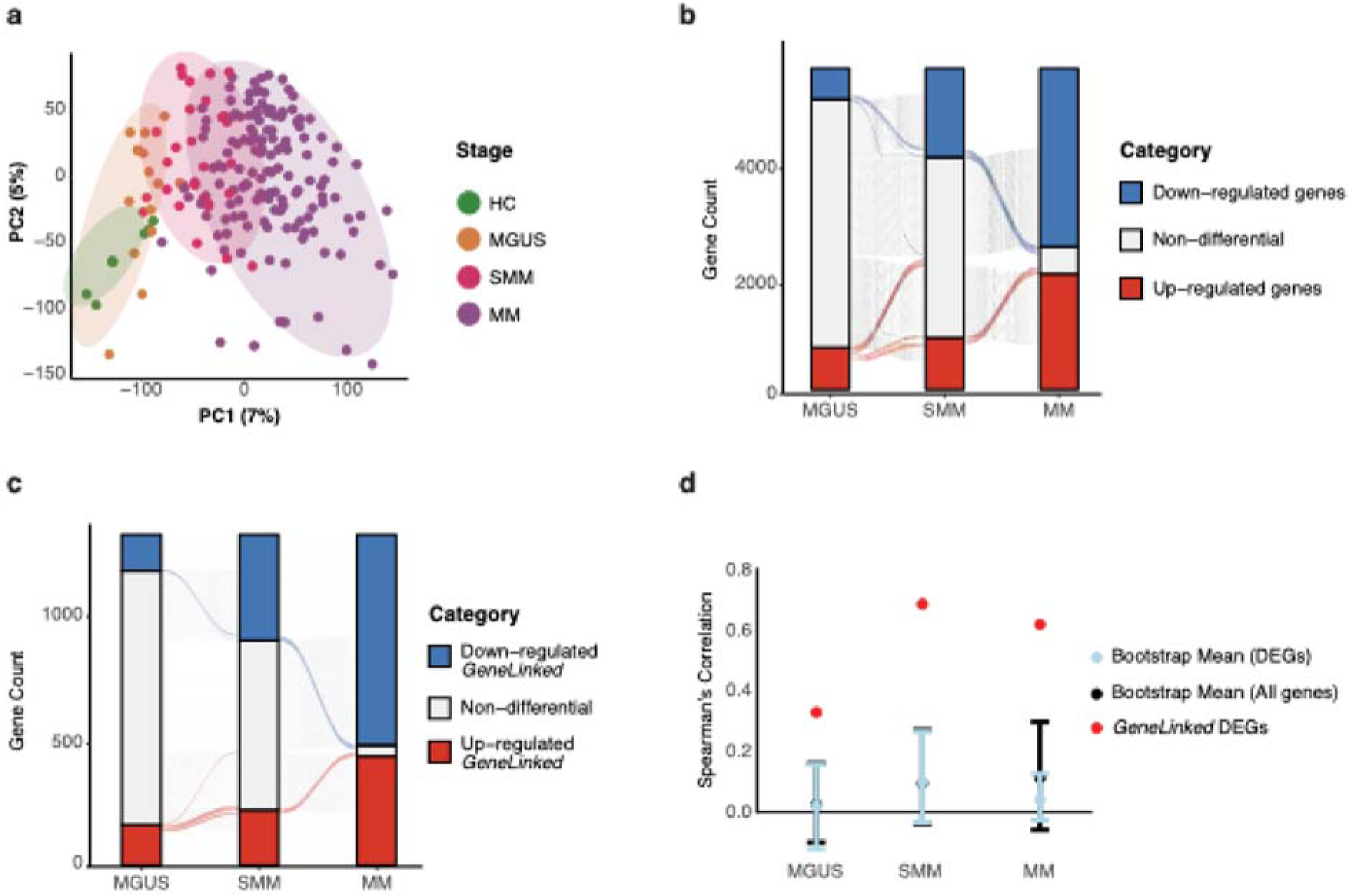
Chromatin accessibility dynamics shape transcriptional programs during myeloma development through increased enhancer-based regulation. a. PCA of normalized RNA-seq gene-expression profiles across all samples (n = 197). Each point represents a single sample colored by disease stage. Ellipses indicate group dispersion. b. Alluvial plot showing significantly DEGs identified by bulk RNA-seq analysis in MGUS vs HC, SMM vs HC, and MM vs HC comparisons. Red indicates upregulated genes, blue indicates downregulated genes, and gray indicates the rest. c. Alluvial plot restricted to the subset of DEGs identified linked to differential chromatin accessibility through significant OCR–gene associations (*LinkedGenes*). Colors indicate genes that are upregulated (red), downregulated (blue), or not differentially expressed (gray) across disease stages. d. Validation of the *LinkedGenes* set in the *<u>validationCohort</u>*^17–19^. For each gene set, the figure shows the mean Spearman correlation between bulk and single-cell expression shifts across HC and disease stages. Red dots represent “DEGs within *LinkedGenes*” set. To control for gene set size, light-blue dots indicate the bootstrap-derived median correlation obtained from randomly sampled DEG sets matched in size to the DEGs within *LinkedGenes* set. Black dots represent the corresponding random samples drawn from the complete set of expressed genes matched in size to the DEGs within *LinkedGenes*. Error bars indicate bootstrap-derived confidence intervals generated through gene-set resampling.

To validate at the transcriptomic level the cumulative accumulation of disease-specific OCRs and differentially expressed genes (DEGs) identified in our bulk discovery cohort, we generated a pseudobulk-based validation cohort (*validationCohort*) using publicly available single-cell datasets^17–19^ spanning HC (n=13) and precursor disease stages (MGUS n=10, SMM n=18, MM n=23) (see Methods). We observed that transcriptomic analysis of the *validationCohort* recapitulated the cumulative transcriptomic changes observed in our bulk RNA seq data (**Supplementary Fig. 2B**). Furthermore, DEGs identified in our bulk discovery cohort showed concordant expression dynamics across disease stages in the *validationCohort* (HC vs MGUS: correlation = 0.33; HC vs SMM = 0.69; HC vs MM: 0.62) (**Supplementary Fig. 2C**).

To investigate regulatory relationships, we linked chromatin accessibility to gene expression using the subset of DARs positively correlated with H3K27ac signal. Promoter OCRs were assigned directly to genes within ±1 kilobase (kb) of the transcription start site (TSS), whereas enhancer–gene relationships for intronic and distal intergenic OCRs were inferred from significant Spearman correlations within a 1 megabase (Mb) TSS-centered window. We identified 13,127 promoter OCRs and 20,246 putative enhancer-associated OCRs (adjusted p < 0.05) (see Methods; **Supplementary Fig. 2D**). For clarity, in what follows, genes identified in at least one OCR–gene pair are referred to as *LinkedGene*, and OCRs involved in at least one OCR–gene pair are referred to as *LinkedOCR*. Notably, among *LinkedGenes* and *LinkedOCRs*, the number of DEG-DAR pairs increased monotonically with disease progression (HC vs MGUS: 194 DEGs, 237 DARs, 246 pairs; HC vs SMM: 475 DEGs, 572 DARs, 617 pairs; HC vs MM: 1,247 DEGs, 1,527 DARs, 1,879 pairs), indicating a stepwise expansion of regulatory rewiring during disease development (**Fig. 2C, Supplementary Fig. 2E, 2F and Supplementary Table 3**).

Importantly, when comparing transcriptional changes between our cohort and the *validationCohort*, *LinkedGenes* showed greater concordance than either all DEGs or randomly selected gene sets, indicating that genes associated with regulatory chromatin changes are more likely to be consistently validated across independent datasets (**Fig. 2D**). Interestingly, analysis of higher-order chromatin organization using Hi-C data from BMPC of 3 patients with MM revealed that OCR–gene pairs are preferentially located within active chromatin compartments (**Supplementary Fig. 2G)** with the upregulated pairs predominantly located within topologically associating domains (TADs) (**Supplementary Fig. 2H).**

In summary, these results demonstrate that progressive chromatin accessibility modulation, orchestrates coordinated transcription, underlying stepwise changes in gene expression during myeloma development.

### Increased enhancer-based regulation during disease development

We next sought to characterize the epigenetic nature of upregulated *LinkedOCRs* across disease stages. Comparison with previously established chromatin-state annotations in MM cells^10^ showed that promoter-associated regulatory activation was already evident in early disease stages and persisted throughout disease development. Notably, upregulated *LinkedOCRs* progressively accumulated in regions originating from low-signal heterochromatic states, increasing from MGUS (20.39%) to SMM (25.43%) and MM (29.53%). Conversely, a substantial fraction of active promoters in disease emerged from poised promoters in nPCs (MGUS: 42.1%; SMM: 36.21%; MM: 26.01%) (**Fig. 3A**).

**Figure 3.**
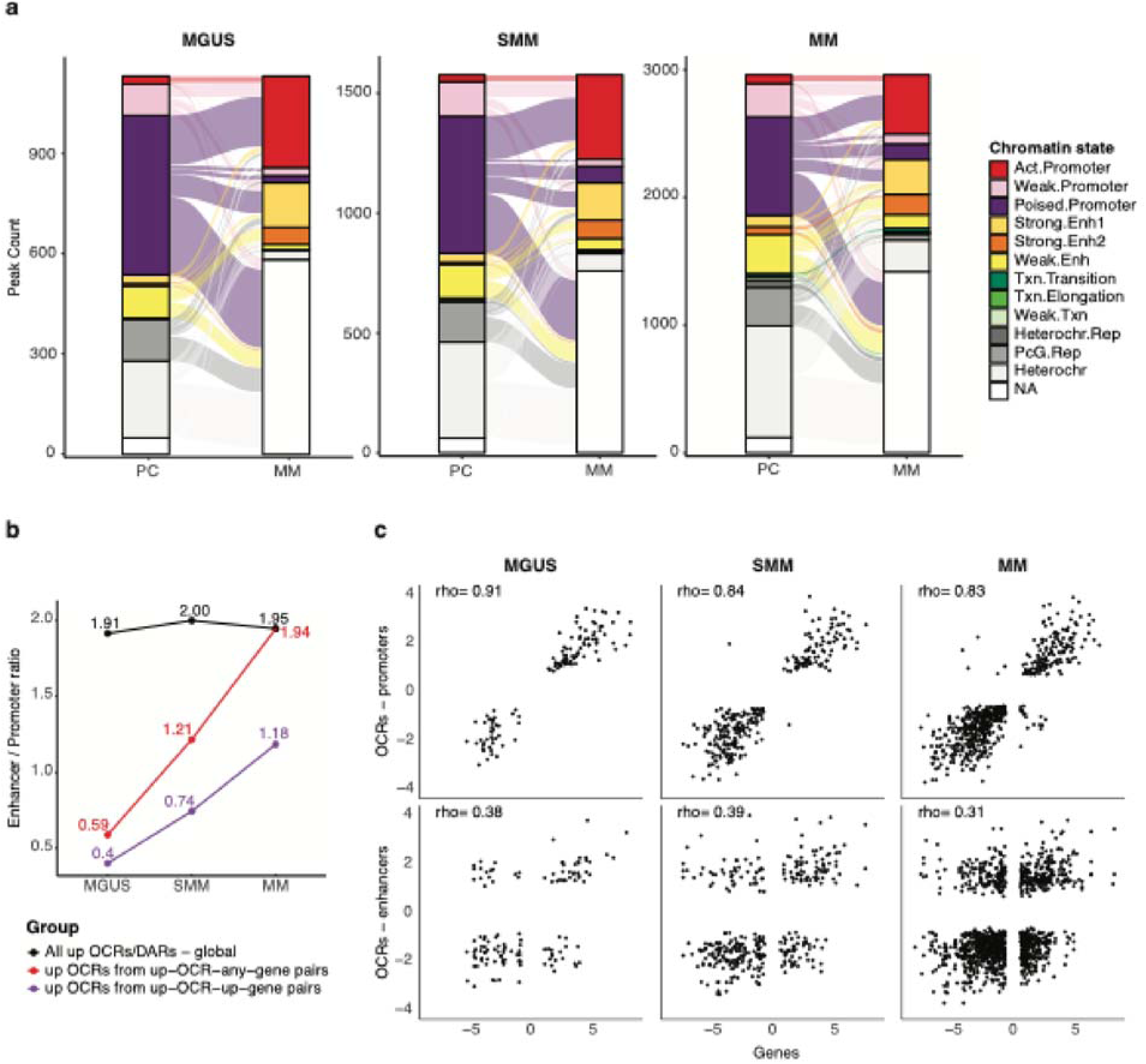
Increased enhancer-based regulation during disease development. a. Stacked bar plots showing chromatin-state annotation of upregulated *LinkedOCRs* across MGUS, SMM, and MM. Chromatin-state annotation was performed separately for *LinkedOCRs* identified in each disease-stage comparison relative to HC, resulting in distinct PC distributions across panels. States include active promoter, weak promoter, poised promoter, strong enhancer 1/2, weak enhancer, transcription-associated states, heterochromatin, polycomb-repressed, and unannotated regions. b. Line plot showing enhancer-to-promoter ratios across MGUS, SMM, and MM. Black line indicates all upregulated DAR OCRs. Red line indicates all upregulated *LinkedOCRs*. Purple line indicates upregulated *LinkedOCRs* paired with upregulated *LinkedGenes*. c. Scatter plots comparing log fold-change values of chromatin accessibility in *LinkedOCRs* (y-axis) vs log fold-change values of gene expression in their associated *LinkedGenes* (x-axis) across disease contrasts. Upper panels correspond to promoter-associated *LinkedOCRs*; lower panels correspond to enhancer-associated *LinkedOCRs*. Each point represents one OCR–gene pair. Spearman correlation coefficients (*rho*) are shown in each panel.

Therefore, we next investigated whether enhancer-driven regulation increased during disease development. Across all regulatory elements, the global enhancer-to-promoter ratio was 1.54. Among upregulated DARs, this ratio was higher in disease relative to HC but remained stable across stages (MGUS: 1.91; SMM: 2.00; MM: 1.95; black line) (**Fig. 3B**), indicating an early global shift toward enhancer-driven regulation. In contrast, when focusing specifically on significantly upregulated *LinkedOCRs*, we observed a clear stage-dependent redistribution from promoter- to enhancer-based regulation (MGUS: 0.40; SMM: 0.742; MM: 1.18; purple line). Yet, and as expected, when comparing shifts in gene expression and chromatin accessibility, promoter-linked OCR–gene pairs showed strong positive correlations, while enhancer-linked pairs displayed weaker correlations (**Fig. 3C**).

Together, these results indicate that promoter-associated transcriptional activation is established early during MM progression and maintained throughout disease development, while enhancer-mediated regulatory activity progressively increases and likely contributes to reinforcing disease-specific transcriptional programs.

### Epigenetically driven regulatory networks activate myeloma-associated pathways

Next, we aimed to identify epigenetically regulated pathways by performing over-representation analysis (ORA) on upregulated *LinkedGenes* (see Methods; **Supplementary Fig. 2F and Supplementary Table 4**). This approach allowed us to prioritize gene sets that are transcriptionally shifted during disease and are associated with epigenetically driven upregulation. We grouped the identified gene sets in seven categories: growth-related pathways, Wnt signaling (Wnt), cellular and microenvironment remodeling, bone-related processes, phosphatidylinositol 3-kinase (PI3K) signaling, cell migration and chemotaxis (**Fig. 4A**). At early stages of the disease (MGUS), we identified migration and chemotaxis pathways as epigenetically regulated, together with early activation of core survival signaling programs such as PI3K and Wnt cascades. Furthermore, as disease progressed to SMM and MM, an increasing number of genes within these pathways became activated (**Fig. 4B**), indicating progressive reinforcement of these signaling programs. In addition, SMM and particularly MM exhibited the emergence of distinct epigenetically regulated programs associated with cellular and microenvironmental remodeling as well as bone-associated processes. These changes are consistent with enhanced tumor–microenvironment interactions and established myeloma bone disease^20,21^. Importantly, similar stage-dependent pathway activity patterns were observed in the *validationCohort* profiles^17–19^ (**Fig. 4C**).

**Figure 4.**
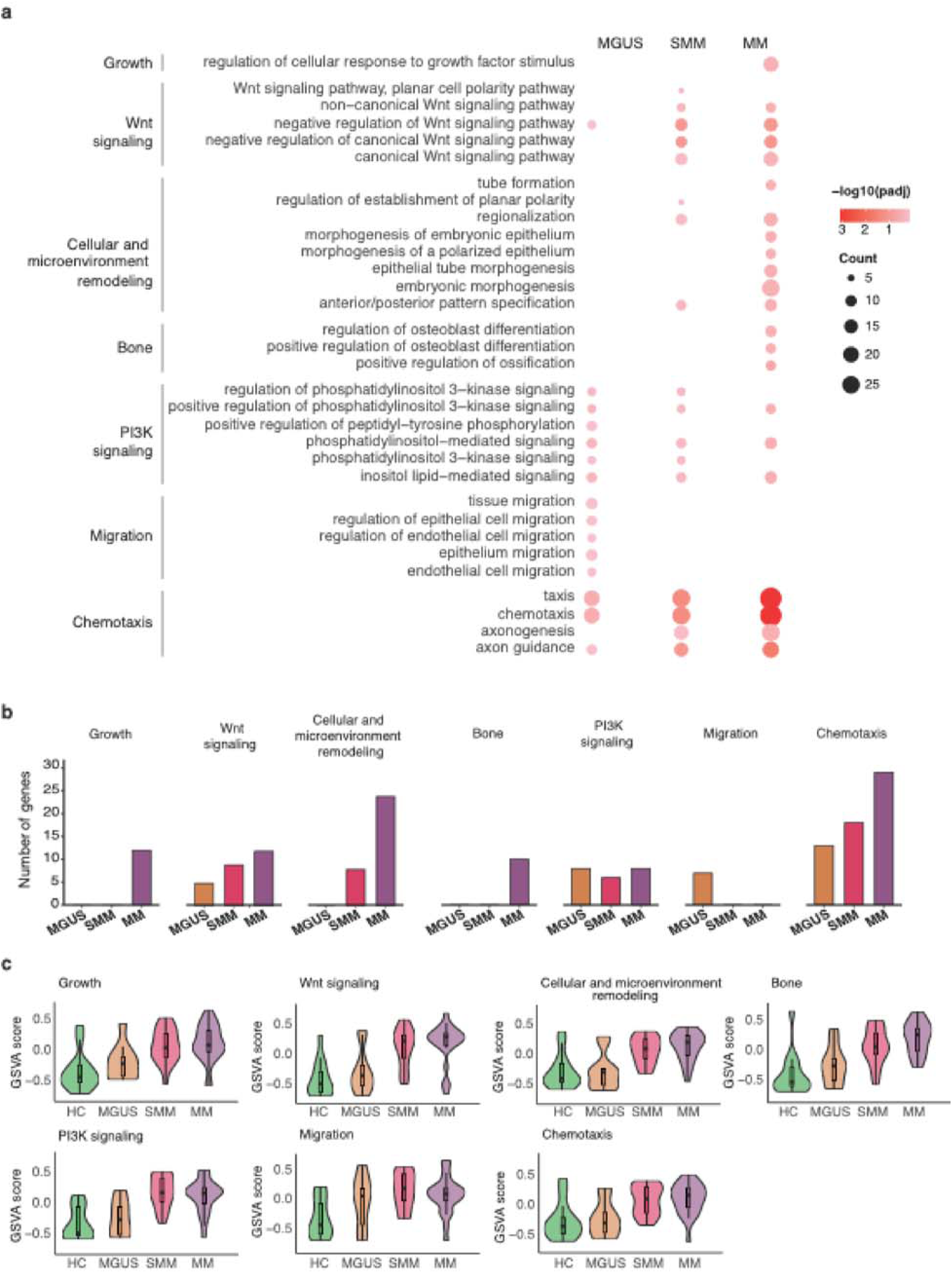
Epigenetically-driven regulatory networks activate myeloma-associated pathways. a. Dot plot showing ORA of upregulated *LinkedGenes* across MGUS, SMM, and MM. Each row represents one enriched pathway term. Dot size indicates the number of genes contributing to enrichment, and color intensity indicates statistical significance (−log10 adjusted P value). Pathways were grouped into functional categories: growth, Wnt signaling, cellular and microenvironment remodeling, bone, PI3K signaling, migration, and chemotaxis. b. Bar plots representing the number of *LinkedGenes* assigned to each pathway category across MGUS, SMM, and MM. Bars are colored by disease stage. c. Violin plots showing GSVA pathway activity scores across HC, MGUS, SMM, and MM using the *validationCohort* datasets^17–19^ for growth, Wnt signaling, cellular and microenvironment remodeling, bone, PI3K signaling, migration, and chemotaxis.

To identify candidate regulators of these pathways, we analyzed TF binding site (TFBS) enrichment within *LinkedOCRs* that progressively gained accessibility across disease stages. This analysis recovered several TFs previously described as essential regulators of MM, including MYC^22,23^, MAF^24–27^, and several members of the IRF family^13,15,28–30^. The number of regulatory interactions associated with these TFs increased from MGUS to MM, consistent with the progressive expansion of accessible chromatin regions across disease stages (**Supplementary Table 5**). Furthermore, these findings suggest that the identified TFs may play a crucial role in the underlying biology of both MM and its precursor conditions, MGUS and SMM. Importantly, this highlights a broad landscape of previously underappreciated TFs that warrant further investigation.

To further support these observations, we computationally inferred TF-associated pathway regulatory potential. For each TF, pathway, and disease stage, candidate target genes were identified, and a corresponding regulon activity score was computed in each of the datasets comprising the *validationCohort* ^17–19^ (see Methods). This analysis confirmed a stage-dependent increase in activity across disease progression. TFs associated with chemotaxis (FOXK2, FOXK1, POU2F2), PI3K (MEF2D, MEF2C, MEF2A, POU2F2), and members of the IRF family showed higher scores at later disease stages, consistent with the regulatory patterns observed in our results (**Fig. 5A**). Of note, these increases were accompanied by substantial inter-patient heterogeneity (**Supplementary Fig. 3A**).

**Figure 5.**
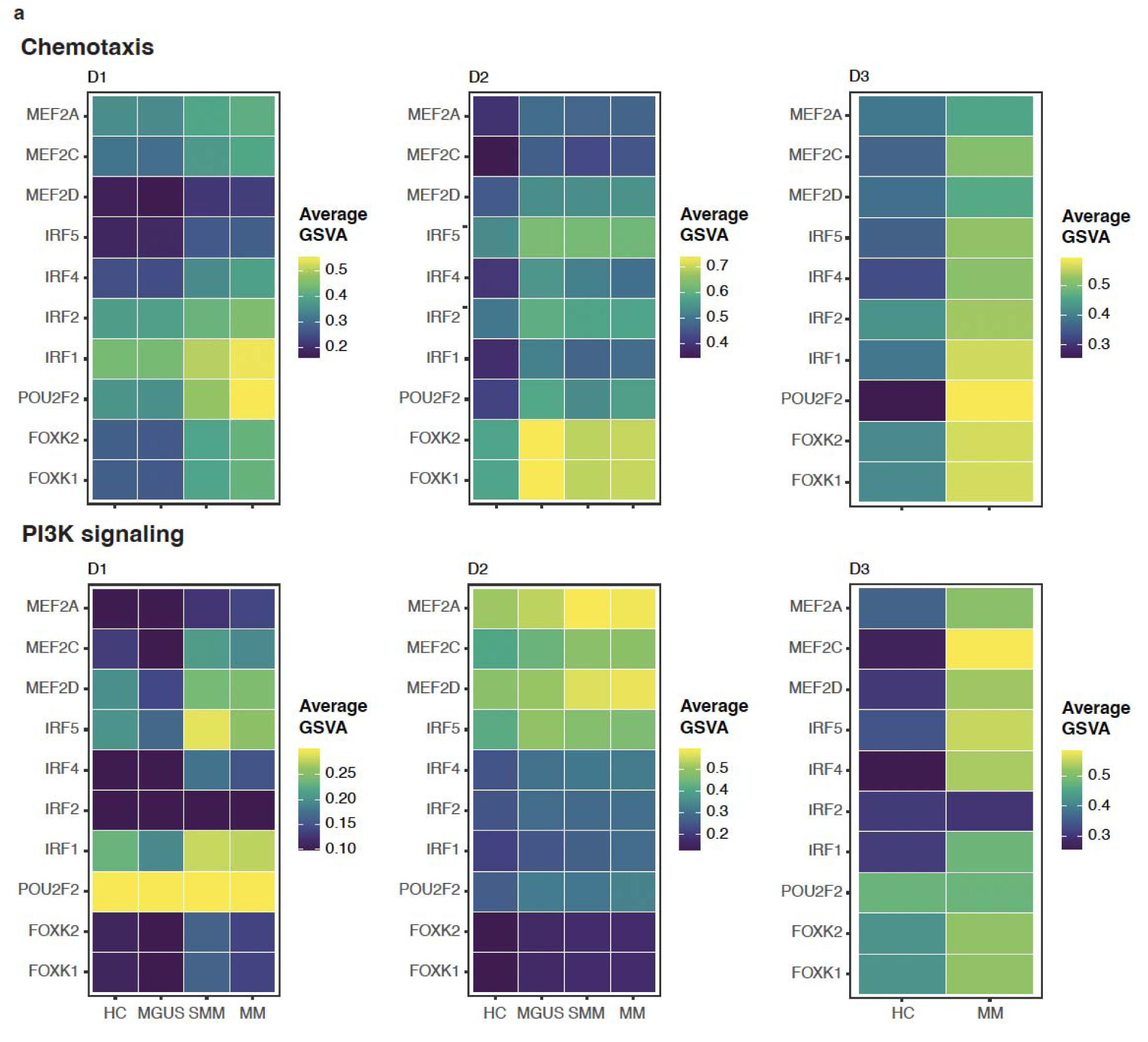
TF-associated pathway regulatory potential validation in public single-cell datasets. a. Heatmaps showing inferred average TF activity scores (rows) across disease stages (columns) from public single-cell datasets (D1: Boiarsky et al., D2: Dang et al., D3: Chen et al.)^17–19^. Separate panels display chemotaxis, and PI3K signaling programs.

Consistent with our previous findings, where CUT&RUN was utilized to define the gene targets of IRF1, IRF2, and IRF4^15^, we observed a significant enrichment of the previously identified targets within the OCR-gene targets of the IRF family defined in our current study (**Supplementary Fig. 4A**). This experimental validation underscores the fidelity of our integrative analysis in capturing functional regulatory interactions.

Together, these results suggest that the identified myeloma-associated transcriptional programs in this study are epigenetically established in precursor states and progressively reinforced through TF regulation during disease development.

### FOXK2 and MEF2D regulate enhancer-mediated transcriptional programs in myeloma

To further validate the functional relevance of disease-associated TFs identified as regulators of OCR–gene pairs, we cross-referenced them with a recent CRISPR/Cas9 screening study performed by our group^15^. Several TFs associated with chemotaxis- and PI3K-related pathways were also identified as essential in this screening (**Supplementary Fig. 5A**). Among these, MEF2D and FOXK2 regulate OCR–gene pairs that persist from MGUS through SMM to MM and expand at later stages (**Fig. 6A, 6B**), suggesting progressive expansion of their regulatory programs. *MEF2D* was expressed similarly in healthy and MGUS PCs, but its expression significantly increased in SMM and MM cells (**Fig. 6C**). Within the MMRF CoMMpass Study cohort^31^ *MEF2D* was also significantly upregulated in patients with high-risk (HR) genetic alterations (**Supplementary Fig. 5B**). Moreover, higher *MEF2D* expression was also associated with significantly poorer Progression-Free Survival (PFS) and Overall Survival (OS) (**Fig. 6D**), suggesting its potential as a prognostic biomarker. FOXK2 showed weaker essentiality in the CRISPR/Cas9 screening than MEF2D (**Supplementary Fig. 5A**). *FOXK2* median expression in SMM and MM cells was higher than in MGUS and healthy PCs (**Fig. 6E**) and in the MMRF CoMMpass Study cohort^31^, the stratification of patients by FOXK2 levels revealed that higher expression correlated significantly with reduced PFS and OS (**Fig. 6F**).

**Figure 6.**
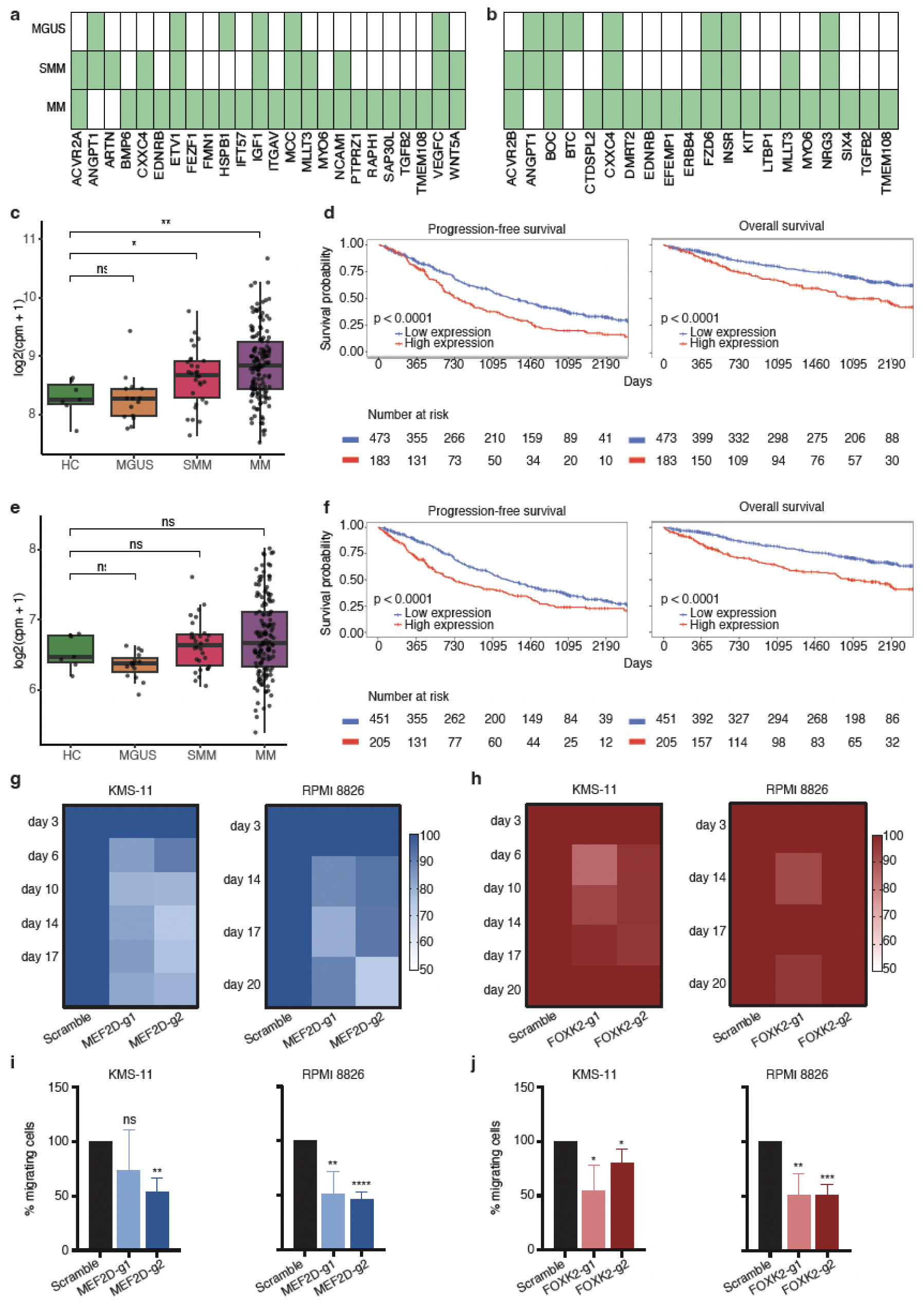
FOXK2 and MEF2D regulate enhancer-mediated transcriptional programs in myeloma. a,b. Representation of significantly associated OCR–gene pairs assigned to MEF2D (a) and FOXK2 (b) across MGUS, SMM, and MM. Connections indicate OCR-linked target genes containing enriched binding motifs for the indicated TF. c. Boxplot showing normalized *MEF2D* RNA expression across HC, MGUS, SMM, and MM samples. Significance is indicated as ns, P < 0.05, P < 0.01. d. Kaplan–Meier curves showing progression-free survival (PFS; left) and overall survival (OS; right) in MM patients from the MMRF CoMMpass cohort^31^. Patients were stratified by high vs low *MEF2D* expression. Two-sided log-rank P values are shown. Numbers at risk are displayed below each plot. e. Boxplot showing normalized *FOXK2* RNA expression across HC, MGUS, SMM, and MM samples. Statistical comparisons were performed as in panel (b). f. Kaplan–Meier curves showing PFS (left) and OS (right) in MM patients from the MMRF CoMMpass cohort^31^ stratified by high vs low *FOXK2* expression. Two-sided log-rank P values are indicated, with numbers at risk shown below the curves. g. Heatmaps showing relative cell viability over time following MEF2D knockout in KMS-11 and RPMI-8226 cells. Rows indicate time points and columns indicate sgRNA conditions. Values are normalized to scramble controls (set to 100%). h. Heatmaps showing relative cell viability over time following FOXK2 knockout in KMS-11 and RPMI-8226 cells. Values were normalized to scramble controls. FOXK2 depletion caused a moderate viability defect with partial recovery at later time points. i. Transwell migration assays following MEF2D knockout in KMS-11 and RPMI-8226 cells. Bar plots show migrating cells expressed as percentage relative to scramble control. Bars represent mean ± SEM from three independent biological replicates. j. Transwell migration assays following FOXK2 knockout in KMS-11 and RPMI-8226 cells. Quantification is shown as percentage of migrating cells relative to scramble control (mean ± SEM; three independent biological replicates).

MEF2D and FOXK2 were inhibited in two MM cell lines (KMS-11-Cas9 and RPMI 8226-Cas9) using two independent single guide RNAs (sgRNAs) per TF. Efficient depletion was confirmed by western blot (**Supplementary Fig. 5C, 5D**), after which cell viability and chemotactic behavior were evaluated. MEF2D inhibition resulted in a marked reduction in MM cell viability (**Fig. 6G**), whereas FOXK2 inhibition caused a moderate decrease that partially recovered over time (**Fig. 6H**). Notably, targeting either TF significantly impaired the ability of MM cells to move through the transwell in response to the chemotactic stimulus (**Fig. 6I, 6J, Supplementary Fig. 5E, 5F**), demonstrating the role of these TFs in the regulation of MM cell chemotaxis.

These findings suggest that inhibiting these TFs in MGUS or SMM cells may affect chemotactic signaling, potentially limiting their ability to locate and reach microenvironments that support their adaptation, survival, and proliferation. In addition, targeting MEF2D may have further implications, as its inhibition was associated with reduced cell viability, which could hinder myeloma clonal expansion.

## Discussion

Understanding how precursor conditions relate to MM at the regulatory level remains a central challenge. Detection of precursor stages enables timely intervention and improved overall survival in MM patients^32,33^. However, the asymptomatic nature of these stages and the low abundance of malignant BMPCs constrain their detection and isolation, as well as the establishment of immortalized cell lines for detailed molecular characterization^2,34,35^. While previous studies have demonstrated widespread epigenetic remodeling in MM, these efforts have largely focused on symptomatic disease state, relapse cases, or specific cytogenetic subgroups, leaving early regulatory events associated with precursor conditions insufficiently characterized^8,36–42^.

Here, by integrating chromatin accessibility, histone acetylation, and gene expression profiles across disease stages we map coordinated epigenetic remodeling landscape underlying the transition from precursor conditions to active MM. Our integrative multi-omic analyses identify a cumulative regulation pattern in which epigenetic alterations are established early in precursor states and reinforced during disease development.

A central finding of this study is that epigenetic remodeling in MM follows a stepwise pattern. Chromatin regulatory elements associated with MM progressively accumulate across disease stages, including regions originating from previously inactive or low-signal chromatin states. This extends prior reports of *de novo* enhancer activation^10^, by demonstrating that such regions are already detectable at early stages and progressively reinforced during disease development, rather than emerging exclusively at symptomatic transformation.

The interplay between epigenetic regulation and transcriptional programs represents a fundamental feature of cancer pathogenesis^43–45^, including in MM^16,46,47^. In line with this, we also demonstrate that chromatin accessibility changes are closely linked to transcriptional alterations through OCR–gene associations that increase during disease development. This expansion supports a model in which early epigenetic alterations contribute to a permissive landscape that is subsequently exploited to reinforce disease-associated transcriptional programs. Notably, our data suggest that promoter-associated transcriptional regulation is established early in precursor conditions and maintained throughout disease progression, whereas enhancer-mediated regulatory activity progressively expands through newly accessible distal regulatory elements, likely reinforcing and diversifying disease-associated transcriptional programs. Similar enhancer-orchestrated regulation has been reported in other malignancies^48–50^

At the functional level, our data indicate that myeloma-associated signaling programs are epigenetically established early and progressively amplified. Pathways related to migration, chemotaxis, and core survival signaling, including PI3K and Wnt cascades, are already epigenetically activated in precursor states, consistent with their roles in PC homing, persistence within the BM, and interactions with the tumor microenvironment^21,51–54^. As the disease advances, these programs expand and are complemented by the emergence of additional pathways linked to cellular and microenvironmental remodeling and bone-associated processes. These findings suggest that early epigenetic changes enable niche engagement and PC survival, while subsequent pathways enhance tumor fitness, increase dependence on tumor microenvironment, and drive disease-specific adaptations. The identification of early epigenetic alterations in MGUS and SMM indicates that regulatory changes precede adverse malignant transformation and may represent potential targets for early intervention.

Our analyses further highlight the contribution of TFs to these stage-specific regulatory dynamics. The enrichment of TF binding motifs within the broader expansion of OCR–gene pairs suggests increased TF occupancy at newly activated regulatory elements. Known MM regulators^55^, including MYC^23^, MAF^24,27^, and members of the IRF family^15,28,30^, show increasing regulatory interactions across disease stages, consistent with their established roles in MM biology. In addition, MEF2D and FOXK2, which have been implicated in normal hematopoiesis and other B-cell associated neoplasms^56,57^, also emerge as candidate regulators of transcriptional programs involved in MM development. Both TFs, show progressive expansion of their associated regulatory targets across disease stages. The association of MEF2D with high-risk genetic alterations and adverse clinical outcomes further supports its potential clinical relevance. Consistent with these observations, similar roles for MEF2D and FOXK2 have been described in other malignancies, where they function as context-dependent regulators of oncogenic transcriptional programs, controlling cell survival, proliferation, and enhancer-mediated gene expression. For instance, MEF2D has been implicated in hematological malignancies, where it participates in enhancer-driven transcriptional regulation^58^. In solid tumors, MEF2D has also shown to promote progression through epigenetic activation, driving epithelial–mesenchymal transition (EMT)^59^, and metastatic potential^60^. Similarly, FOXK2 regulates oncogenic transcriptional programs in different cancers, modulating gene expression networks linked to proliferation, metabolism, and survival through chromatin-associated mechanisms^61,62^.

Functional perturbation experiments demonstrate that MEF2D contributes to cell viability and that both MEF2D and FOXK2 influence chemotactic behavior, linking epigenetic regulatory network expansion to phenotypic outputs. These findings suggest that the association MEF2D and FOXK2 with key myeloma-related pathways, such as chemotaxis and survival signaling, may act as early regulators that are progressively engaged during disease development. Importantly, although functional validation was performed in MM cell lines, the multi-omic epigenetically informed analyses spanning the full spectrum of disease stages, together with the validation in external single-cell datasets^17–19^, support the functional relevance of these inferred networks. These observations raise the possibility that targeting such TF-associated programs might modulate pathways potentially involved in malignant transformation.

Despite the reproducibility of our findings across independent datasets, several limitations should be noted. Although our cohort represents a comprehensive multi-epigenomic resource spanning MM disease stages, its cross-sectional design precludes direct assessment of longitudinal progression within individual patients. Most importantly, functional validation was performed in established MM cell lines, which may not fully capture the biology of early disease, reflecting the limited availability of experimental models for MGUS and SMM.

In summary, our study provides a comprehensive, stage-associated framework of epigenetic and transcriptional remodeling across myeloma development. We show that epigenetic alterations are established early in precursor states and progressively reinforced through coordinated promoter-associated, enhancer-mediated, and TF-driven regulatory programs. Moreover, the progressive nature of regulatory rewiring highlights opportunities for stage-specific therapeutic strategies, targeting early promoter-driven programs, or later expanding enhancer-associated networks. These findings provide a foundation for understanding early disease states and support further investigation into whether regulatory elements or their upstream factors may serve as potential targets for disease stratification or therapeutic intervention. The identification of candidate regulators, such as MEF2D, provide a basis for future studies aimed at evaluating epigenetically informed therapeutic strategies and biomarker development.

## Methods

### Human sample collection

Human BM samples were obtained from healthy donors (n=7) and newly diagnosed MGUS (n=15), SMM (n=30), and MM (n=145) patients through the Biobank of the University of Navarra, following approved standard operating procedures and ethical approval from the local Research Ethics Committee. Our cohort included individuals carrying recurrent myeloma-associated cytogenetic lesions detected by fluorescent in situ hybridization (FISH), including translocation t(4;14), translocation t(14;16), chromosome 17p deletion (del(17p)), chromosome 1p deletion (del(1p)), and gain and/or amplification of chromosome 1q (referred to herein as 1q aberrations) (**Fig. 1B**). Patients included both sexes with median age of 71 years.

### FACS sorting

Mononuclear cells were obtained from BM samples after Ficoll-Isopaque density centrifugation and cryopreserved for future use. After thawing, CD138+ PCs were isolated by flow cytometry cell sorting using FACSAria II. Sorted plasma cells were collected in a lysis buffer for bulk RNA-Seq sequencing or 1X PBS, 0.05% BSA for ATAC-Seq and ChIP-Seq.

### Fluorescent *in situ* hybridization (FISH)

FISH was performed using a panel targeting regions of common cytogenetic abnormalities in MM. A minimum of 100 interphase cells were dropped onto a glass slide and dried briefly before fixation *in situ*. The panel consists of two probes, 1q21 and 1p32, to detect gains, as well as deletions of chromosome 1, respectively; a locus specific probe, 17p13, to detect TP53 deletion and a break-apart probe to detect rearrangements of IGH (14q32). Rearrangements of IGH were further investigated with IGH/MAF and IGH/FGFR3 dual color probes.

Our cohort encompassed patients with key myeloma-associated genetic lesions identified by FISH, including t(4;14), t(14;16), del(17p), del(1p), and gain and/or amplification of 1q, hereafter collectively termed 1q aberrations (**Fig. 1B**), while 77 samples showed no detectable alterations.

### ATAC-seq data

Briefly, 10,000 cells were sorted in 1X PBS, 0.05% BSA and pelleted by centrifugation at 500 rcf for 5 min at 4 ÅãC with low acceleration and brake settings in a pre-cooled swinging-bucket rotor centrifuge. All the supernatant but 5 μL was removed. Next, 25 μL of transposase mix (15 μL of 2x TD buffer (Illumina); 1 μL of TDE1 (Illumina), 0.25 μL of 5% digitonin (Promega); 8.75 μL of nuclease-free water) were added to the cells and the pellet was disrupted by pipetting. Transposition reactions were incubated at 37 ÅãC for 30 min in an Eppendorf ThermoMixer with shaking at 450 rpm. Reactions were stopped at 4 ÅãC for 5 min. In order to release tagmented DNA, samples were incubated at 40 ÅãC for 30 min with 5 μL of clean up buffer (900 mM NaCl (Sigma), 30 mM EDTA (Millipore), 2 μL of 5% SDS (Millipore) and 2 μL of Proteinase K (NEB)). DNA was purified using a 2X SPRI beads cleanup kit (Agencourt AMPure XP, Beckman Coulter). In order to determine the total number of PCR cycles needed for library amplification, two sequential PCRs were performed using KAPA HiFi DNA Polymerase and customized Nextera PCR indexing primers (IDT) (Buenrostro et al, 2013). The conditions of the first PCR were: 5 min at 72 ÅãC and 2 min at 98 ÅãC followed by 9 cycles of 20 secs at 9 ÅãC, 30 secs at 63 ÅãC, and 1 min at 72 ÅãC. Depending on the library concentration in this first PCR, a second PCR (2 min at 98 ÅãC followed by 4 to 6 cycles of 20 secs at 98 ÅãC, 30 secs at 63 ÅãC, and 1 min at 72 ÅãC) was performed aiming for a library concentration in the range of 2 to 10 ng/μL. PCR products were purified using a 2X SPRI beads cleanup. Libraries were quantified and their size profiles examined as described above. Sequencing was carried out in an Illumina NextSeq 500 using paired-end, dual-index sequencing (Rd1: 38 cycles; Rd2: 38 cycles; i7: 8 cycles; i5: 8 cycles).

### ChIP-seq data

ChIP-seq of H3K27ac histone mark data were generated as described (http://www.blueprintepigenome.eu/index.cfm?p=7BF8A4B6-F4FE-861A-2AD57A08D63D0B58), following the high-quality standards of Blueprint Consortium (EU contribution to the International Human Epigenome Consortium). Catalog number of antibodies (Diagenode) used for H3K27ac: C15410196/pAb-196-050. In order to minimize any technical problems (i.e. Ab specificity), all these antibodies were broadly tested and validated from different perspectives before being applied for ChIP-Seq experiments. All the data concerning the validated Blueprint antibodies as well as established SOPs are publicly available on Blueprint-Epigenome website (http://www.blueprintepigenome.eu/index.cfm?p=D6F8811F-DACF-7979-CEAC0B9034C28037).

### RNA-seq data

Publicly available RNA-seq data from our previous work GSE302052^15^.

### ATAC-seq quantification and normalization

Reads’ quality was assessed with FastQC software (v0.11.8) (Babraham Institute). Low quality, short reads, and Illumina adapters were trimmed using Trimmomatic (v0.38). Reads were then mapped to the hg38 reference genome (GRCh38 Ensembl annotation release 103) using Bowtie2 (v2.3.4.2). Unmapped, unknown, random, ChrM mapping, duplicate reads, and reads matching to blacklist (the blacklist includes problematic regions of the genome^63^) were filtered out using samtools view -bh -F 1028 (SAMtools v1.9), and Picard Tools (v2.18.17) (Picard MarkDuplicates, http://broadinstitute.github.io/picard). Paired-end reads were used to determine those regions with significant reads coverage in contrast to the random background (regions known as peaks or OCRs) using MACS2 (v2.1.0.2015) (function callpeak) with the default parameters (https://github.com/taoliu/MACS). A minimum peak size of 20 base pairs was set as a threshold. Fraction of reads in peaks (FRIP) score was calculated to assess enrichment of reads in peak regions. FRIP was calculated as the ratio of reads in peaks to the total number of reads. To get the consensus OCRs for each myeloma population, we found the genome regions which are covered by at least 2 samples for a specific myeloma population of the sets of peaks and then we merged the nearby peaks with a minimum gap width of 30 base pairs. To get the consensus peaks in the whole population, we merged all of the common and unique regions in the genome from the different myeloma subpopulations and then merged the nearby peaks with a min gap width of 30. These processes were done using the rtracklayer and GenomicRanges r-packages. Using the whole consensus OCRs, we computed the count matrix using the *multicov* function from bedtools. Those OCRs without a minimum of 10 counts for at least one sample and 15 total counts were excluded, retaining 142,136 OCRs. To address potential batch effects in the ATAC-seq data, batch correction was performed using the *removeBatchEffect* function from the limma package. Here, the FRIP score was used as a proxy for batch information. The raw ATAC-seq counts were log-transformed and a model matrix was constructed using the disease state, aberrations, and sex of the samples to ensure that biological variability was retained during correction.

### RNA-seq quantification and normalization

Reads’ quality was assessed with FastQC software (v0.11.8) (Babraham Institute). Low quality, short reads, and Illumina adapters were trimmed using Trimmomatic (v0.38). Reads were then mapped to the December 2013 Homo sapiens high coverage assembly GRCh38 from the Genome Reference Consortium GRCh38 (hg38) using STAR47 and counted for each gene using htseq-count with –m intersection-nonempty option and a GTF file from GRCh38 Ensembl annotation release 103, getting the information for a total of 60,666 genes, after removing Ig genes. Genes with a minimum read count of 1 in all population replicates and an associated ensemble ID were retained (24,256 genes). Counts were normalized under TMM normalization and voom transformation from edgeR r-package (v4.0.14).

### Estimating the proportion of normal plasma cells: nPCs proxy score

Despite the genetic diversity within our cohort, variability could be driven by the relative proportions of malignant and nPCs in each patient. Following the assumption that HC samples consist entirely of nPCs, and MM samples entirely of malignant PCs, the proportions of these cell types in MGUS and SMM may vary. To identify changes specific to MGUS and SMM progression, it is essential to account for the proportion of malignant cells in each sample. Since a direct estimate is unavailable, we inferred a nPCs gene activity score (hereafter referred to as % nPCs) for each individual using their transcriptional profile and a reference gene set comprising flow cytometry markers commonly used to distinguish nPCs (CD19, CD27, CD38, CD45 and CD81). The gene activity score was computed using Gene Set Variation Analysis (GSVA) enrichment score method in R, applied to RNA-seq gene expression data. As expected, the estimated %nPCs declines progressively from MGUS to MM, reflecting the increasing burden of mPCs with disease progression (**Fig. S1A**).

### Differential analysis

Two linear models were applied to both ATAC-seq and RNA-seq datasets for differential accessibility and differential expression analyses, respectively:

A. ã*=stage+aberration+sex*
B. ã*=stage+aberration+sex+%nPCs*

Model A was applied to contrasts between HC and malignant stages (MGUS, SMM, or MM), with the union of all differential results capturing global changes across disease progression. Model B specifically focused on the transitions from HC to MGUS and HC to SMM, incorporating %nPCs to identify stage-specific features, while minimizing bias from variations in nPCs proportions in precursor stages. Consensus features for downstream analysis, were defined as the intersection between global changes identified by Model A and % nPCs-corrected features identified from Model B. For the HC to MM transition, only Model A was applied.

Differential analyses were performed using limma (v3.58.1) with voom transformation. DARs were selected using logFC > |0.58| and adjusted p < 0.01, whereas DEGs were selected using logFC > |0.58| and adjusted p < 0.05. A more stringent p-value threshold was applied to ATAC-seq data to prioritize robust regulatory changes within the larger and more variable chromatin accessibility feature space.

### ChIP-seq quantification and normalization

Raw sequencing reads quality was assessed using FastQC (v0.11.8). Reads were aligned to the human reference genome (GRCh38/hg38) using Bowtie2 (v2.3.4.2) with the following parameters: --very-sensitive, --no-discordant, and --no-mixed, allowing a maximum fragment length of 1000 bp. The resulting alignments were processed with SAMtools (v1.9) for BAM format conversion and coordinate sorting.

To mitigate amplification biases, library complexity was estimated and PCR duplicates were marked using Picard (v2.18.17). Unmapped reads and marked PCR duplicates were subsequently excluded from downstream analyses (SAMtools flag -F 1028). Further filtering was applied to remove genomic artifacts: ENCODE blacklist regions for hg38 were excluded using BEDTools (v2.27.1) and reads mapped to the mitochondrial chromosome (chrM) or unplaced contigs (chrUn) were discarded.

For genomic visualization, coverage profiles in BigWig format were generated from the filtered BAM files using the bamCoverage tool from deepTools (v3.2.0). The signal was normalized to RPKM (Reads Per Kilobase per Million mapped reads) using a bin size of 20 bp and an effective genome size of 3Gbp. Enriched region identification (peak calling) was performed using MACS2 (v2.1.0) with default parameters for narrow peaks. Finally, the signal-to-noise ratio and overall immunoprecipitation quality were estimated by calculating the Fraction of Reads in Peaks (FRiP) score utilizing BEDTools and SAMtools.

A consensus peak set was established by merging the overlapping peaks from individual samples (format converted to Simplified Annotation Format, SAF). Quantitative assessment of the ChIP-seq signal was performed using the *featureCounts* function from the Rsubread package in R. Read counting was executed in a non-strand-specific manner, strictly considering uniquely mapped reads to prevent artificial signal inflation in repetitive genomic regions. Only read pairs with both ends successfully and appropriately aligned were included in the final count matrix. The resulting raw count matrix was subsequently exported for downstream analysis.

Those peaks without a minimum of 10 counts for at least one sample and 15 total counts were excluded, retaining 93,674 peaks. To address potential batch effects in the ChIP-seq data, batch correction was performed using the *removeBatchEffect* function from the limma package. Here, the FRIP score was used as a proxy for batch information. The raw ChIP-seq counts were log-transformed and a model matrix was constructed using the disease state, aberrations, and sex of the samples to ensure that biological variability was retained during correction.

### ChIP-seq integration

To refine the selection of DARs, we integrated ChIP-seq data to identify true active regulatory regions. Since H3K27ac profiling was unavailable for HC samples, we could not perform differential acetylation analysis in a manner analogous to ATAC-seq. Instead, we assessed the relationship between chromatin accessibility and histone acetylation by identifying overlapping regions between ATAC-seq and ChIP-seq peaks and exploring their association using Spearman’s correlation (adjusted p < 0.05; minimum overlap of 6 bp). This approach identified 69,140 acetylated regions that positively correlated with 77,361 accessible regions, yielding a total of 77,616 overlapping pairs. The overlapping pairs were subsequently used to filter DARs, ensuring the selection of regions with active regulatory potential.

### Variance component analysis

Chromatin accessibility variance across ATAC-seq peaks and gene expression variance across RNA-seq data were quantified using the variancePartition linear mixed-model approach, which partitions the total variation attributable to multiple biological and technical factors by fitting a mixed model to each feature while jointly accounting for all other covariates. Normalized accessibility measures (voom log2 counts) were modeled with disease stage, sex, recurrent cytogenetic lesions as random effects, and continuous % nPCs scores as a fixed effect in the formula ∼ (1|Stage) + (1|Sex) + (1|1q aberration) + (1|del(1p)) + (1|del(17p)) + (1|t(14;16)) + (1|t(4;14)) + % nPCs. Pairwise dependencies among metadata variables were evaluated using canonical correlation analysis to identify collinearity. For each peak, the variance fractions attributable to each factor and the residual were extracted using *fitExtractVarPartModel* function and summarized genome-wide by sorting variables by median variance explained, with results visualized via violin plots to illustrate the relative contribution of each source of variation (**Fig. S1D, S2A**).

### Chromatin states assignment to OCRs

We defined chromatin states of our OCRs based on the previously published data^10^. First, we defined consensus chromatin states of peaks across multiple samples. For 4 MM samples presented in the paper we retained the same category of chromatin states in peaks, if it was present in at least 3 or more samples. For 3 PC samples we followed the same logic and retained the same category of chromatin states in peaks, if it was present in at least 2 or more samples. In case of not reaching a consensus (multiple chromatin states in one peak), we labeled that peak as not applicable (NA)). Next, we performed overlaps of MM and PC peaks and our OCRs using *findOverlaps* function from GenomicRanges package (v1.54.1).

### OCRs association with target genes

To investigate the features coordinated between ATAC-seq and RNA-seq, we focused on OCRs mapping to a promoter, intronic or distal intergenic regions. For promoter OCRs, we directly annotated OCRs to genes based on TSS distance (+/- 1kb), thus identifying 13,127 OCRs mapping to promoter regions. For intronic and distal intergenic regions (considered here as putative enhancer regions), we linked OCRs to genes following the hypothesis that Spearman correlation between chromatin accessibility of an enhancer and the expression of a given gene denotes a likely functional connection. Our data revealed 20,246 significantly associated OCRs mapping enhancer regions within a 1Mb window around TSS (+/- 500kb from TSS) [adjusted p < 0.05].

### Gene-set enrichment analysis (GSEA)

To functionally characterize transcriptional changes during disease progression from MGUS to SMM to MM, we performed GSEA on differential expression results for each disease stage compared with healthy controls. Genes were ranked by log fold change. Analyses were conducted in R (v4.2.2) using the clusterProfiler package (v4.6.2). Gene Ontology Biological Process (GO-BP) terms were tested with the *gseGO* function (keyType = “ENSEMBL”, minGSSize = 10, maxGSSize = 500), using gene annotations from org.Hs.eg.db (v3.16.0). Multiple testing correction was applied using the Benjamini–Hochberg procedure, with significance defined at adjusted p < 0.05. Ensembl IDs were converted to gene symbols using the *setReadable* function. Results were summarized and visualized using the binary cut clustering method implemented by the simplifyEnrichment package (v1.8.0).

### Overrepresentation analysis (ORA)

ORA was performed to identify GO-BP pathways enriched among upregulated genes associated with active OCRs in each disease stage. Gene sets were selected based on two criteria: (i) significant enrichment in the transcriptomic analysis (GSEA; adjusted p < 0.01; positive normalized enrichment score), and (ii) significant overrepresentation of genes associated with upregulated OCRs in each disease comparison. The gene universe consisted of all upregulated genes detected in at least one disease-versus-healthy comparison. Enrichment was assessed using the *enricher* function from clusterProfiler (v4.6.2) with Benjamini–Hochberg correction. Pathways with adjusted p < 0.05 were considered significant. Results were summarized using Jaccard index and biological interpretations to group pathways into broader functional categories.

### OCRs association with TFs

TF–OCR associations were computed using *matchMotifs* function with JASPAR position weight matrices (PWMs) to scan OCR sequences for motif presence, generating a binary matrix of motif presence per region.

### Single-cell RNA-seq public data collection and processing

All datasets from Boiarsky et al.^17^, and Dang et al.^18^ were publicly available and accessed from their respective repositories: dbGaP (phs001323.v3.p1), the European Genome-Phenome Archive (EGA; EGAS00001006694), respectively. Chen et al.^19^ was obtained directly from the corresponding author upon request. Raw reads from scRNA-seq FASTQ files were aligned to the human reference genome (hg38) using CellRanger (v7.1.0), creating a gene expression matrix for each sample consisting of UMI counts per gene, per barcoded cell. Using Seurat (v.5.0.3), raw count of each sample was further processed into a Seurat object for preprocessing and QC. To retain high-quality cells, filtering criteria were applied. Cells were excluded if they exhibited a log10(Genes per UMI) ratio below the 80th percentile, mitochondrial gene content exceeding 15%, or if they fell within the top or bottom 10% of total RNA counts and detected features, based on their respective distributions. Additionally, potential doublets were identified and removed using the *scDblFinder* function. Immunoglobulin genes and sex genes (*XIST* and *RPS4Y1*) were removed. Each sample was normalized using *NormalizeData* function. Cell-cycling score was computed per cell using features associated with S and G2M phases. Single-cell RNA-seq data were first aggregated into pseudobulk profiles. For each sample, gene expression values were averaged across all cells using the *AverageExpression* function from the Seurat R package (v.5.0.3). Matrices from all datasets were merged and normalized using counts per million (CPM). Principal component analysis (PCA) was applied to the normalized pseudobulk expression matrix to investigate sources of variance. Associations between the top 50 principal components and sample metadata were evaluated. Kruskal–Wallis tests were used to assess relationships with categorical variables (e.g., stage, dataset source, sex), while Spearman’s correlations were computed for continuous variables (e.g., library size, mitochondrial and ribosomal content). Results were summarized as –log (p) values and correlation coefficients to visualize the influence of technical and biological factors, including potential batch effects. To mitigate such effects, batch correction was applied using the *ComBat* algorithm. Differential expression analysis was performed using the *limma* package with the *voom* transformation. A linear model was fit with the design *∼ stage + batch + sex*, and contrasts were defined to compare each disease stage to healthy controls. For the contrasts between HC and MGUS and between HC and SMM, the model included the proportion of %nPCs as an additional covariate. Significance was determined using empirical Bayes moderation (|log2FC| > 0.58; adjusted p < 0.05).

### Bulk signature validation in scRNA-seq public data

To evaluate consistency between pseudobulk and bulk RNA-seq results, logFC were correlated using Spearman’s method for genes derived from OCR–gene pairs (n = 179, 432, and 1,094 for MGUS, SMM, and MM, respectively). In addition, 1,000 iterations of bootstrapped random gene sets (matched in number per contrast) were sampled from the bulk DEGs or all significantly DEGs, and Spearman correlations were computed. Confidence intervals were calculated to assess variability and determine significance relative to the null distribution.

### Pathway GSVA at a single-cell level

Pathway activity was quantified using Gene Set Variation Analysis (GSVA) applied to batch-corrected pseudobulk expression data. Gene sets were defined by taking the union of stage-specific signatures (MGUS, SMM, and MM) for each pathway (e.g., chemotaxis), ensuring consistent representation across disease progression. GSVA was performed using the *gsva* function to compute enrichment scores per sample, providing a measure of relative pathway activity across disease stages.

### TF-targets score per pathway at a single-cell level

To quantify transcription factor (TF) target activity at single-cell resolution, gene set enrichment scoring was performed separately for each dataset using the scGSVA framework. For each pathway of interest, TF-specific target gene sets were defined based on previously TF–target lists (e.g., chemotaxis-associated targets), and only genes present in the corresponding expression matrix were retained. Enrichment scores were then computed on raw RNA count data using the *scgsva* function with the single-sample GSEA (ssgsea) method, generating per-cell activity scores for each TF-target gene set. The resulting scores were added to the Seurat object metadata and used for downstream comparisons across patients and disease stages.

### Hi-C

Human BM samples were collected from 3 newly diagnosed MM patients. BMPCs were isolated as described in the previous methods section followed by the FISH. *In situ* Hi-C was performed based on the previously described protocol^64^. Briefly, 2 million crosslinked cells per sample were digested with DpnII, filled in with bio-dCTP, and ligated. Nuclei were treated with RNAseA and proteinase K, and DNA was purified, precipitated, and resuspended in Tris buffer. Purified DNA was sonicated, mixed with magnetic streptavidin T1 beads, and subjected to end-repair, A-tailing, and ligation with Illumina adaptors. Libraries were amplified by PCR, purified with AMPure XP beads and quantified by Qubit. Libraries were sequenced on HiSeq 2500.

### Public Hi-C data

Public Hi-C data from healthy PCs were obtained from EGAS00001004763^65^. PCs were isolated from tonsils of male pediatric donors (2–12 years) undergoing tonsillectomy at Clínica Universidad de Navarra (Pamplona, Spain).

### Hi-C preprocessing and analysis

Hi-C reads were aligned to the reference genome GRCh38 using *bwa mem* (v0.7.19) with “-SP5M”. Invalid data, including PCR duplicates and read pairs mapping to the same restriction fragment, were removed using *pairtools* (v0.3.0). *runHiC* (v0.8.7) and *cooler* (v0.10.2) were used to construct contact matrices at various resolutions. Raw Hi-C matrices were corrected using a modified matrix balancing method to account for CNV effects and other systematic biases including mappability, GC content, and restriction enzyme sites, all processed via *Neoloopfinder* (v0.4.3.post2).

PC1 was calculated, and A-/B-compartments identified at a resolution of 100 kilobase pair (kbp) using the *cooltools* (v0.7.1) *call-compartment* function. TADs were identified at 50 kbp resolution using the *OnTAD* (v8.312). Chromatin loops were identified at 10-, and 25 kbp resolution on the basis of interaction probabilities > 0.95 and then merged using *peakachu* (v2.3). For 10 kbp loops, we extended anchors by 5 kbp when searching for associated TSSs to define loop-associated genes; for 25 kbp-resolution loops, no extension was applied.

TADs and loops were compared between HC and MM samples using Jaccard similarity. TADs were considered identical based on reciprocal overlap (≥80%) using findOverlapsOfPeaks from ChIPpeakAnno. Chromatin loops were considered the same when the midpoint of each anchor in one loop was within ±1 bin of the corresponding anchor midpoint in the other loop.

To quantify “recovery” of OCR–gene pairs in 3D chromatin structures, we tested whether each OCR and its paired gene TSS were supported by shared TAD membership and/or loop connectivity. For TAD support, OCRs and TSSs were independently assigned to TADs by genomic overlap, and an OCR–gene pair was considered recovered if both elements mapped within the same TAD (reciprocal domain membership). For loop support, loop anchors were represented as genomic intervals and an OCR–gene pair was considered recovered if the OCR overlapped one loop anchor and the paired gene TSS overlapped the corresponding opposite anchor of the same loop. Recovery was summarized as the fraction of OCR–gene pairs supported by TADs, loops, or both.

### Cell culture

KMS-11 and RPMI 8226 MM cell lines were engineered to stably express Cas9, and maintained in RPMI-1640 supplemented with 20% FBS, 2% HEPES, 1% penicillin–streptomycin, and L-glutamine (Gibco). HEK293T cells were used for lentiviral production and titration in DMEM containing 10% FBS, 20 mM HEPES, and 100 U/ml penicillin–streptomycin. All cells were incubated at 37 °C with 5% CO, authenticated by STR profiling, and regularly tested for Mycoplasma contamination using the MycoAlert kit (Lonza).

### CRISPR/Cas9 inhibition of TFs

Two sgRNAs per TF were chosen from the CRISPR–Cas9 library and packaged into lentiviral vectors, which were used to transduce Cas9-expressing KMS-11 and RPMI 8226 cell lines. Because the sgRNA plasmid encoded Blue Fluorescent Protein (BFP), cell viability was monitored for 20 days by flow cytometry through tracking of the BFP cell population. At seven days post-transduction (dpt), a subset of cells was collected to evaluate protein expression levels by Western blot. The lentiviral particles containing the sgRNAs were produced, titrated and transduced into MM cells as previously described^15^. All the experiments were performed in three biological replicates.

### Chemotaxis analysis by transwell assay

At 10 dpt, 1×10^5^ cells in 100 μl serum-free medium were seeded on the transwell (Costar) placed in a 24-well plate containing 500 μl of 20% FBS culture medium. Seven hours later, cells were fixed, stained and images were taken as previously described^15^. All the experiments were performed in three biological replicates.

### Western blot

Protein detection by Western Blot was carried out as previously described^15^, using the specific primary antibodies for the detection of FOXK2 (ABE1828, Sigma-Aldrich) and MEF2D (ab282731, abcam). All the experiments were performed in three biological replicates.

### Survival analyses

RNA-seq data from 656 patients gathered in the IA18 release of the Multiple Myeloma Research Foundation (MMRF) CoMMpass Study dataset was used^31^. Patients were stratified into high and low expression groups according to MEF2D or FOXK2 expression levels. PFS and OS were evaluated using Kaplan–Meier survival curves with two-sided log-rank tests and univariate Cox proportional hazards regression models, as previously described^15^.

## Data availability

All ATAC-seq, H3K27ac ChIP-seq and HiC data generated in this study will be publicly available as of the date of peer-reviewed. Raw bulk-RNAseq data has been deposited at GSE302052. The public scRNA-seq datasets from Boiarsky et al., and Dang et al. used for validation are available on dbGaP (phs001323.v3.p1) and the European Genome-Phenome Archive (EGA; EGAS00001006694).

