## Supplementary Figures for "Mapping Disease Transitions from Premalignant, Asymptomatic to Advanced Myeloma through Integrative Epigenomic and Transcriptional Analyses"

**SUPP. FIGURE 1**

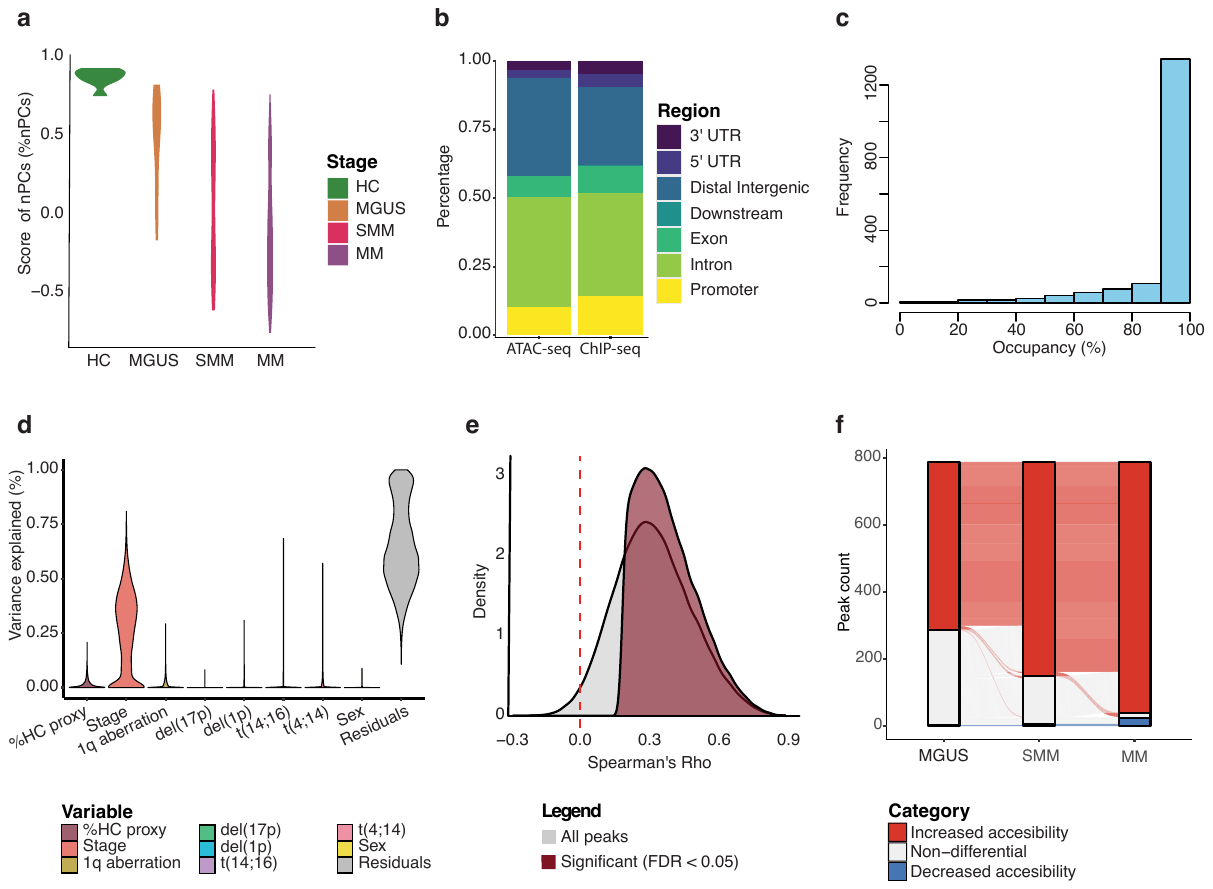

**Figure S1. Additional characterization of progressive chromatin remodeling during multiple myeloma development.**

a. Violin plots representing the distribution of the %nPCs proxy score across HC, MGUS, SMM, and MM. This proxy was derived as described in Methods and used to estimate the contribution of non-malignant PC contamination to bulk chromatin accessibility profiles. The y-axis represents the normalized % nPCs proxy score.

b. Stacked bar plots showing the relative genomic distribution of all identified OCRs (ATAC-seq, n=142,136 peaks) and H3K27ac active regions (ChIP-seq, n=93,674 peaks) across annotated genomic elements. Peak annotations were assigned to promoter, exon, intron, downstream (≤300 bp), distal intergenic, 5′ UTR, and 3′ UTR regions. The y-axis represents the fraction of total peaks assigned to each annotation category.

c. Histogram showing the proportion of each previously described *de novo* active regulatory regions in MM^10^ covered by the identified OCRs in our bulk ATAC-seq. The x-axis indicates the proportion of each reference region covered by overlapping OCR signal (0–100%), and the y-axis represents the number of regions within each occupancy bin.

d. Violin plots showing the distribution of the proportion of variance explained for chromatin accessibility signal across DARs by clinical and molecular covariates. Variables tested include percentage healthy-cell proxy (%nPCs proxy), disease stage, recurrent cytogenetic alterations [1q aberration, del(17p), del(1p), t(14;16), t(4;14)], sex, and residual unexplained variance. The y-axis indicates the fraction of total variance explained per OCR.

e. Density plot showing the distribution of Spearman’s rank correlation coefficients (*rho*) between ATAC-seq accessibility signal and H3K27ac ChIP-seq signal across regions with a minimum overlap of 6 bp (n = 77,361 overlapping peaks). The gray distribution represents all overlapping regions, whereas the burgundy distribution indicates regions with statistically significant positive correlations after multiple testing correction (FDR < 0.05). The vertical dashed red line marks rho = 0. The x-axis represents Spearman’s *rho*, and the y-axis indicates density.

f. Bar plot showing the total number of *de novo* active regions overlapping with DARs identified in pairwise comparisons between HC and each disease stage (MGUS vs HC, SMM vs HC, MM vs HC). Red segments indicate regions with increased accessibility, blue segments indicate regions with decreased accessibility, and gray segments represent non-differential OCRs.

**SUPP. FIGURE 2**

**Figure S2. Additional analyses supporting progressive chromatin–transcriptional coupling during MM evolution.**

a. Violin plots showing the distribution of the proportion of variance explained for gene-expression levels across DEGs by clinical and biological variables. Covariates included percentage nPCs proxy (% nPCs proxy), disease stage, recurrent cytogenetic alterations (1q aberration, del(17p), del(1p), t(14;16), t(4;14)), sex, and residual unexplained variance. The y-axis indicates the fraction of total variance explained per gene.

b. Alluvial plot showing the total number of DEGs identified in pairwise comparisons between HC and each disease stage (MGUS vs HC, SMM vs HC, MM vs HC) in the single-cell *validationCohort*^17–19^. Red segments indicate upregulated genes, blue segments indicate downregulated genes, and gray segments represent non-differential genes.

c. Scatter plots showing the correlation of differential gene-expression logFC derived from *validationCohort*^17–19^ (x-axis) and our bulk RNA-seq (y-axis) for MGUS vs HC (left), SMM vs HC (middle), and MM vs HC (right). Each dot represents one gene identified as differentially expressed. Spearman’s rank correlation coefficient (*rho*) and two-sided P values are indicated in each panel.

d. Schematic representation of the analytical strategy used to identify OCR–gene regulatory pairs. Step 1: Detection of all significant OCR–gene associations linking promoter- or enhancer-associated OCRs to putative target genes. Step 2: Selection of disease-relevant OCR–gene pairs by retaining associations in which *LinkedGenes* were DEGs and *LinkedOCRs* were DARs. Red lines indicate OCR–gene regulatory links. Colored boxes denote promoter (P), enhancer (E), transcription start site (TSS), target gene body, and disease-associated elements.

e. Alluvial plot showing the number and directionality of OCR–gene *Linkedpairs* across MGUS vs HC, SMM vs HC, and MM vs HC comparisons. Separate bars are shown for OCRs and linked genes within each contrast. Red indicates coordinately upregulated pairs, blue indicates coordinately downregulated pairs, and gray indicates non-differential pairs.

f. UpSet plot summarizing intersections among significant *LinkedGenes* identified in MGUS vs HC, SMM vs HC, and MM vs HC comparisons, stratified by direction of change (upregulated or downregulated). Horizontal bars indicate total set size for each comparison, whereas vertical bars indicate the number of genes shared across the indicated combinations.

g. Alluvial plot showing assignment of OCR–gene pairs to A (active) or B (inactive) chromatin compartments using Hi-C maps from nPCs (PC) and MM samples (MM1–MM3). The y-axis indicates the number of OCR–gene interactions. Most OCR–gene pairs remained within A compartments, indicating that disease-associated regulatory rewiring occurs predominantly within active three-dimensional genome domains.

h. Bar plot showing the percentage of upregulated OCR–gene pairs per disease contrast localized within TADs chromatin compartments based on Hi-C data from nPCs (PC) and three MM samples (MM1–MM3). Bars represent the proportion of OCR–gene interactions assigned to each sample-specific compartment map. Values above bars indicate percentages.

**SUPP. FIGURE 3**

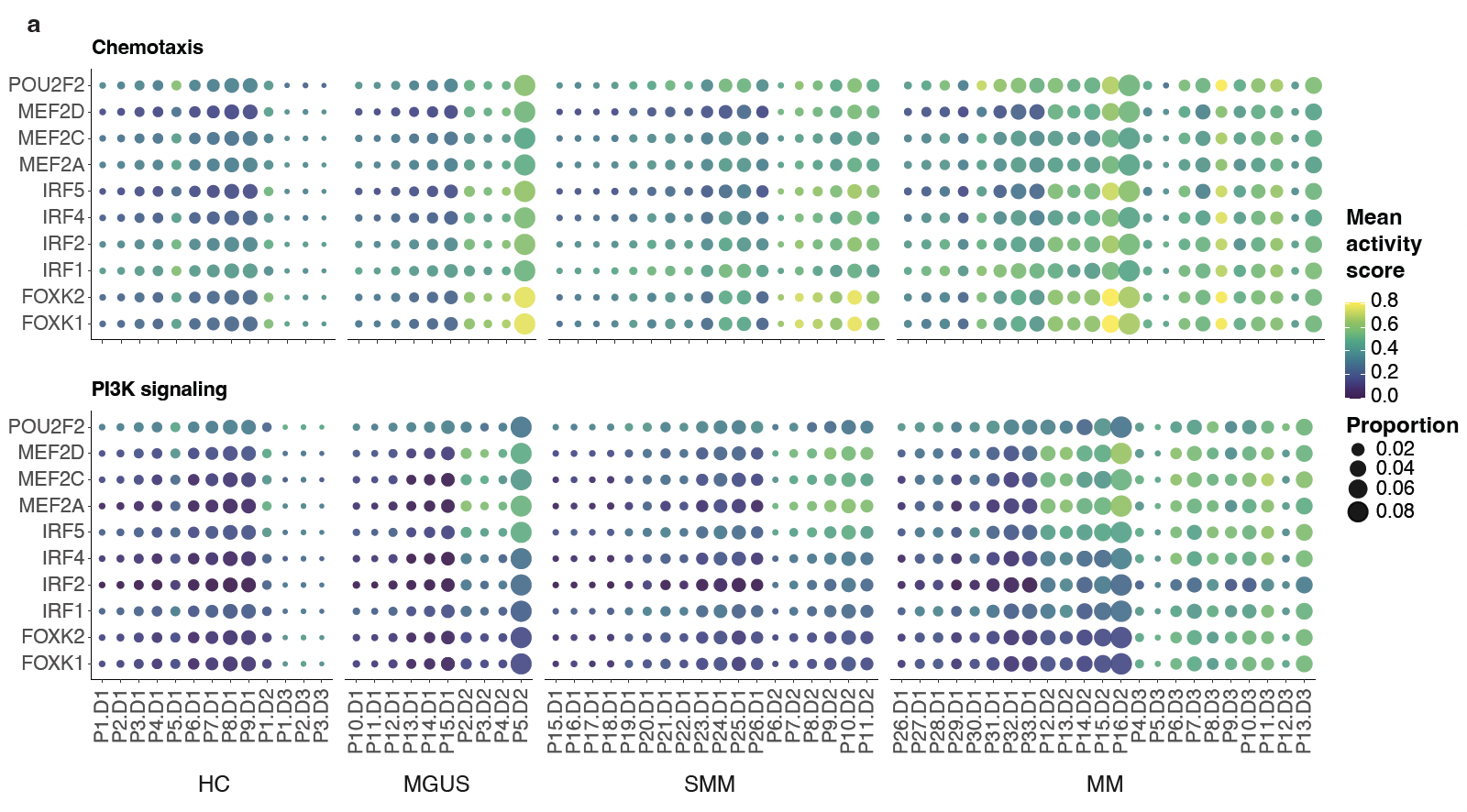

**Figure S3. TF-associated pathway regulatory potential validation in public single-cell datasets.**

a. Dot plot showing inferred average TF activity scores for IRF family, MEF family, FOXK1, FOXK2 and POU2F2 across HC, MGUS, SMM, and MM patients from public single-cells datasets. Separate panels display chemotaxis, and PI3K signaling programs. Each dot represents one patient/sample-level cell subset. Dot size indicates the proportion of cells with detectable activity, and color denotes mean activity score.

**SUPP. FIGURE 4**

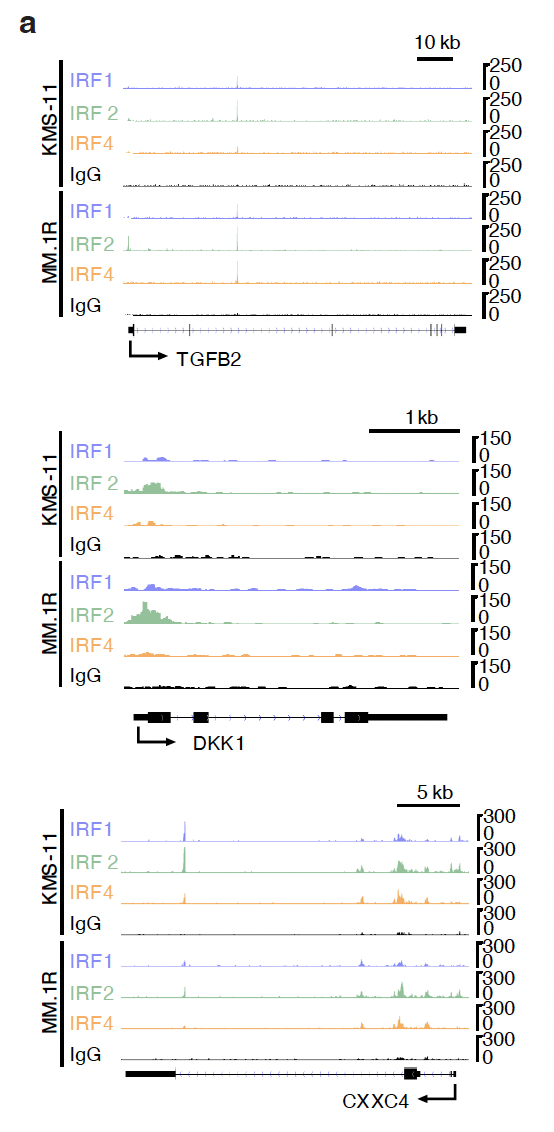

**Figure S4. Independent CUT&RUN validation of IRF-family regulatory targets associated with myeloma-related pathways.**

a. Representative normalized CUT&RUN signal tracks from our previously published dataset^15^ for IRF1, IRF2, IRF4, and IgG control at the *TGFB2*, *DKK1*, and *CXXC4* loci in KMS-11 and MM.1R MM cell lines. Scale bars denote genomic distance (10 kb for *TGFB2*, 1 kb for *DKK1*, and 5 kb for *CXXC4*). Enrichment of IRF-family binding over background (IgG) at these loci supports their classification as candidate IRF-regulated targets identified through OCR–gene linkage analyses in our study.

**SUPP. FIGURE 5**

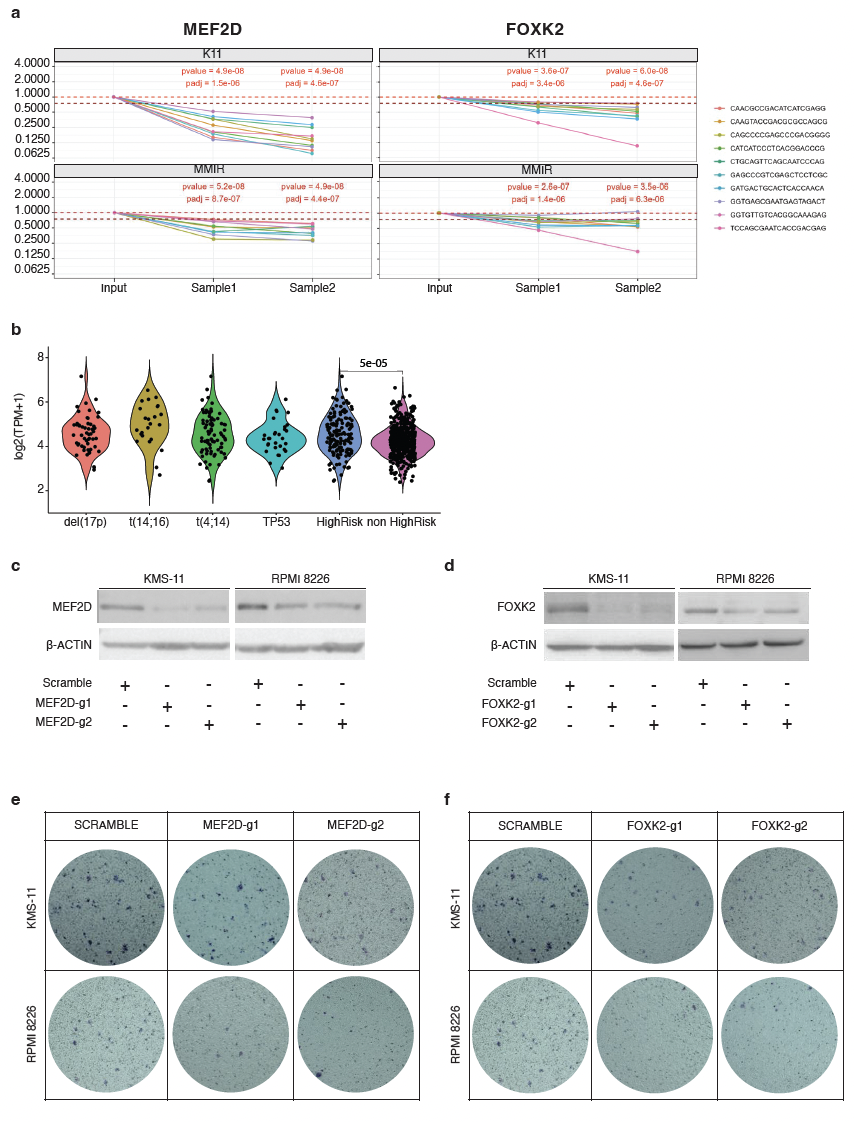

**Figure S5.**

a. Line plots showing relative fitness scores following CRISPR/Cas9-mediated targeting of MEF2D (left) or FOXK2 (right) using 10 independent sgRNAs per gene in two MM cell lines (KMS-11and MM.1R). Each colored line represents one sgRNA. Fitness was measured at day 0, day 14 and day 21. The dashed red horizontal line marks a fitness score of 1, and the dark red dashed line marks a score of 0.25 to facilitate visualization of stronger depletion effects. Nominal P values and multiple-testing adjusted P values (padj) comparing sgRNA depletion over time are indicated within each panel.

b. Violin plots showing normalized *MEF2D* expression across MM molecular subgroups from the MMRF CoMMpass cohort^31^ defined by recurrent alterations, including del(1p), t(14;16), t(4;14) and TP53 alterations, high-risk, and non-high-risk categories. Each dot represents one patient sample. Statistical is indicated in the plot.

c. Western Blot analysis of MEF2D upon its knockdown in two MM cell lines (KMS-11 and RPMI 8226) using two different sgRNAs (MEF2D-sg1 and MEF2D-sg2).

d. Western Blot analysis of FOXK2 upon its knockdown in two MM cell lines (KMS-11 and RPMI 8226) using two different sgRNAs (FOXK2-sg1 and FOXK2-sg2).

e. Representative bright-field images of crystal violet–stained migrated KMS-11 and RPMI-8226 cells collected from the lower surface of transwell membranes after CRISPR/Cas9-mediated MEF2D knockout using two independent sgRNAs (MEF2D-sg1 and MEF2D-sg2). Scramble sgRNA was used as control.

f. Representative bright-field images of crystal violet–stained migrated KMS-11 and RPMI-8226 cells collected from the lower surface of transwell membranes after CRISPR/Cas9-mediated FOXK2 knockout using two independent sgRNAs (FOXK2-sg1 and FOXK2-sg2). Scramble sgRNA was used as control.
